# Molecular archaeology: individual human DNA and infection profiles from 100 year old mosquitoes

**DOI:** 10.1101/2023.05.19.541455

**Authors:** Florence Crombe, Sanjay C Nagi, Elsa VE Tomás, J. Derek Charlwood, Martin J Donnelly, David Weetman

## Abstract

Natural history collections represent a vast biological and genetic resource, which has remained largely untapped. We revisit mosquito collections of varying ages; some made by early tropical medicine pioneers over 100 years ago. By applying species-specific primers, a panel of forensic short tandem repeat (STR) markers, and *Plasmodium* diagnostics, we were able to obtain the unique genetic profile and *Plasmodium* infection status of the individual from whom each mosquito obtained their final bloodmeal. We show evidence of long-dead mosquitoes feeding on multiple individuals in the same gonotrophic cycle, and how the human hosts are likely to have been infected with malaria at the time of blood-feeding. This approach may be used to track the host-fidelity of vectors over an evolutionary time scale.

## Introduction

Archived collections held by natural history museums and research institutes may play an important role in understanding species invasions, responses to climate change, and variation in biodiversity (1–4). Genetic examination of such samples has illuminated ancient evolutionary relationships between parasites and host species (5,6). In vector-borne disease biology, avian tissue from museum collections has been invaluable in tracing the origins of the parasite *Plasmodium relictum*, which has devastated Hawaii’s native avifauna (7) and understand the relationships between extant and eradicated *P. vivax* in *Europe* from historical medical slides.

Insect disease vectors evolve rapidly to overcome human interventions such as insecticide treated nets either through evolution of physiological resistance (8,9) or putative shifts in feeding behaviour (10). Within a single mosquito species, feeding behaviour and host preference can vary dramatically (11,12), resulting in marked heterogeneity in the distribution of mosquito bites across human populations (13). For example, pregnant women often receive a disproportionate number of bites (14). This differential attractiveness can result in marked differences in infection prevalence and disease burdens across a population.

Understanding what drives heterogeneity in transmission risk is increasingly important in the malaria eradication era, as falling endemicity may result in high-burden hotspots of disease that could be targeted with appropriate control (15). Examining host-fidelity of vectors over long time-scales gives us the ability to study the evolution of transmission risk for both anthroponotic and zoonotic diseases.

The bloodmeals of recently-caught mosquitoes have been used to characterise intraspecific variation in vertebrate host selection (12,16), changes to biting behaviour following insecticide-treated bednet introduction (17), forensic profiling of murder victims (18) and to study human*-Aedes* vector contacts to estimate dengue transmission networks (19). In this study we demonstrate that by using up to 100-year-old mosquito specimens, we were able to identify both biting preferences and *Plasmodium* infection status of the person from whom the mosquito obtained their final bloodmeal.

## Methods

### Mosquito dissection and PCR methods for species identification and blood-meal typing

Wild caught specimens were obtained from five collectors; 1. Dr. J.W.S. Macfie was collected in Accra, Gold Coast (present day Ghana), 1915; 2. Mrs. S.L.M. Connal, Lagos, Nigeria, 1915; 3. Prof. Harold Townson, Coastal Kenya, 1979; 4. Dr Eveline Klinkenberg, Accra, Ghana 2004; 5. Dr Derek Charlwood, Cambodia 2012 (Table 1). Blood-fed lab specimens were also used to assess how soon post-blood feeding specimens would have needed to be preserved to permit successful blood meal identification. These specimens were *Anopheles gambiae* from a colony maintained at the Liverpool School of Tropical Medicine, which were fed on a human volunteer and killed at 5 different time points; 0, 12, 24, 36 and 48 hours post blood feeding.

**Table 1:** Summarized description of the samples used for allelic profile identification.

| Sample no. | Origin | Collector | Extraction | Species<br>morphological/<br>molecular ID | *Plasmodium*<br>TaqMan |
| --- | --- | --- | --- | --- | --- |
| Aedes_2012_1 | Cambodia | J. D.<br>Charlwood | Qiagen kit | *Aedes* spp. | - |
| Aedes_2012_2 | Cambodia | J. D.<br>Charlwood | Livak | *Aedes* spp. | - |
| Aedes_2012_3 | Cambodia | J. D.<br>Charlwood | Qiagen kit | *Aedes* spp. | - |
| Odumasy_2006_1 | Odumasy,<br>Ghana | E. Klinkenberg | Livak | *An. gambiae* | - |
| Accra_2004_1 | Accra | E. Klinkenberg | Livak | *An. gambiae* s.l | - |
| Jimbo_1979_1 | Jimbo, Kenya | H. Townson | Qiagen kit | *An. quadriannulatus* | - |
| Jimbo_1979_2 | Jimbo, Kenya | H. Townson | Qiagen kit | *An. quadriannulatus* | + |
| Jimbo_1979_3 | Jimbo, Kenya | H. Townson | Qiagen kit | *An. quadriannulatus* | - |
| Jimbo_1979_4 | Jimbo, Kenya | H. Townson | Qiagen kit | *An. quadriannulatus* | - |
| Jego_1979_5 | Jego, Kenya | H. Townson | Livak | *An. quadriannulatus* | + |
| Accra_1915_1 | Gold Coast<br>Accra, Ghana | J.W.S. Macfie | Qiagen kit | *An. gambiae* | - |
| Accra_1915_2 | Gold Coast<br>Accra, Ghana | J.W.S. Macfie | Qiagen kit | *An. gambiae* | - |
| Accra_1915_3 | Gold Coast<br>Accra, Ghana | J.W.S. Macfie | Livak | *Mansonia* spp. | - |
| Yaba_1915_4 | Yaba, Lagos,<br>Nigeria | S.L.M. Connal | Qiagen kit | *Mansonia* spp. | + |
| Yaba_1915_5 | Yaba, Lagos,<br>Nigeria | S.L.M. Connal | Qiagen kit | *Mansonia* spp. | - |

Wild-caught specimens were morphologically typed to genus level and DNA extracted from mosquitoes which, based on their darkened and distended abdomens, appeared to have bloodfed shortly before their collection. Mosquito heads and thoraces were separated from the abdomens of the collected blood-fed mosquitoes to allow differentiation of malaria parasite detection in salivary glands (infectious sporozoite stage) from the bloodmeal (infected, but not infectious gametocyte stage). DNA was extracted from the abdomens using either a Livak (20) or Qiaquick (Qiagen, Gmbh, Hilden) method. The vertebrate origin of the blood meal was determined using a mitochondrial *Cytb* diagnostic PCR method (21). For mosquitoes morphologically-identified as within the *Anopheles gambiae* species complex, standard molecular species identification assays were performed to identify the species (22,23). Additional confirmatory sequencing of the ribosomal DNA intergenic spacer region (polymorphism within which is targeted by one of the standard species diagnostics) (21) was conducted for a number of specimens (Table 1).

Human blood meals were analysed by amplifying 10 STR (microsatellite) loci (i.e. CSFPO, D3S1358, D7S820, D8S1179, D13S317, D16S539, D18S51, TH01, TPOX and FGA) and the Amelogenin (AMEL) gender-identification locus (24,25) using the Type-it Microsatellite PCR kit (Qiagen Gmbh, Hilden). The three multiplex PCRs and their respective primer pairs are given in Supplementary Table 1. Briefly, amplifications were performed in reaction volumes of 25 μL using a reaction mix containing 1x Type-it Multiplex PCR Master Mix (Qiagen, Gmbh, Hilden), 0.2 μM of each forward and reverse primer pair, except for locus D7S820 which had a final primer concentration of 0.4μM. For recent samples, 1ul of template DNA was used, whereas 6ul was used for historic samples (>20 years) and the time-course experiments. PCR conditions were 95°C for 5 min, followed by 32 cycles at 95°C for 30 s, 60°C for 1.5 min, 72°C for 30 s, with a final extension at 60°C for 30 min. Amplified fragments were genotyped on an Applied Biosystems 3700 sequencer by Macrogen Europe (Amsterdam, The Netherlands) and Source Biosciences (Nottingham, UK). In order to obtain the correct allele number from peak height and size, 6 human samples were sent to Promega (Southampton, UK) and analysed using the PowerPlex16 panel as standards. Subsequent analysis was conducted using GeneMapper v.5 software (Applied Biosystems: Thermo-Fisher). Blood meals were screened for *Plasmodium falciparum* using a TaqMan quantitative PCR assay (26).

## Results and Discussion

Specimen collection dates ranged from 1915 to 2012 and extended geographically from Africa to Southeast Asia (Table 1). All mosquitoes for which species-specific blood meal identification was possible had fed on a human. In all but one case it was possible to identify the gender of the person on which the mosquito took her last meal (Table 2). We tested the accuracy of the panel by application to a blood sample from Dr. Charlwood. The resultant genotypes all matched perfectly those in two of the three specimens he collected (Aedes_2012_2 and 3; Table 2).

**Table 2:** Allelic profiles obtained from the abdomen of blood-fed mosquitoes after DNA extraction. NI: STR loci not identified due to unsuccessful amplification.

| Sample no. | STR loci |  |  |  |  |  |  |  |  |  |  |
| --- | --- | --- | --- | --- | --- | --- | --- | --- | --- | --- | --- |
|  | D7S820 | TH01 | TPOX | D16S539 | D8S1179 | D3S1358 | D18S51 | FGA | D13S317 | CSFP10 | AM |
| Aedes_2012_1 | 11/12 | 7/9.3 | 9/11 | 10/12 | 12/14 | 17/18 | 12/12 | 23/23 | 13/13 | 11/12 | X/Y |
| Aedes_2012_2 | 8/9 | 9/9 | 8/11 | 9/11 | 12/13 | 16/17 | 16/19 | 20/32.2 | 12/13 | 12/12 | X/X |
| Aedes_2012_3 | 11/12 | 7/9.3 | 9/11 | 10/12 | 12/14 | 17/18 | 12/12 | 23/23 | 13/13 | 11/12 | X/Y |
| Odumasy_2006_1 | 8/11 | 6/7/8 | 6/9/11 | 10/11/13 | 14/15/17 | 15/16 | 16/19 | 23/24/25 | 11/12/13 | 12/12 | X/Y |
| Accra_2004_1 | 8/12 | 7/7 | 8/9 | 9/10 | 15/15 | 14/16 | 17/19 | 21/23 | 13/13 | 12/12 | X/X |
| Jimbo_1979_1 | 10/11 | 8/9 | 9/10 | 9/11 | 12/15 | 15/16 | 13/18 | 23/23 | 10/12 | 8/9 | X/Y |
| Jimbo_1979_2 | 10/11/13 | 8/9 | 8/9/10 | 9/11/12 | 12/14/15/16 | 15/16/17 | 13/15/18/19 | 21/23 | 10/12 | 8/9/10 | X/Y |
| Jimbo_1979_3 | 10/12 | 7/8 | 8/12 | 11/12 | 11/14 | 15/16 | 18/18 | 20/25 | 11/12 | 6/12 | X/Y |
| Jimbo_1979_4 | 8/8 | 7/8 | 8/11 | 9/11 | 13/14 | 16/16 | 16/17 | 22/25 | 11/12 | 8/10 | X/Y |
| Jego_1979_5 | 10/10 | 9/9 | 8/11 | 9/13 | 12/13 | 14/15 | 16/17 | 23/27 | 12/12 | 12/12 | X/Y |
| Accra_1915_1 | 8/10 | 7/9.3 | 8/11 | 8/10 | 14/15 | 14/17 | 14/17 | 21/28 | 11/15 | 11/11 | X/Y |
| Accra_1915_2 | 10/11 | 7/9.3 | 9/9 | 10/11 | 14/14 | 17/18 | 16/17 | 23/23 | 11/11 | 8/12 | X/Y |
| Accra_1915_3 | 10/11 | 7/9.3 | 9/9 | 10/11 | 14/14 | 17/18 | 16/17 | 23/23 | 11/11 | 8/12 | X/Y |
| Yaba_1915_4 | NI | 7/8 | 6/8 | 10/11 | 12/15 | 16/17 | 12/15 | 22/24 | 12/12 | 8/8 | X/Y |
| Yaba_1915_5 | NI | 7/9 | 8/11 | 9/9 | NI | NI | NI | NI | NI | NI | NI |

The analysis revealed two clear cases where mosquitoes had fed on more than one individual, indicated by more than two alleles detected per locus across multiple markers (Odumasy_2006_1 and Jimbo_1979_1; Table 2). The accuracy of the older sample’s profile was checked by a second independent genotyping test and provided an identical result at each locus Typically, female mosquitoes feed only once to repletion during each period of oogenesis, suggesting that these mosquitoes may have been disturbed mid-feed and sought an alternative host. In laboratory studies, we demonstrated that it was only possible to identify the blood meal origin up to 24-36 hours post-feeding (Figure S1), and it is therefore unlikely that the blood is from an earlier egg-laying cycle, as the duration of the gonotrophic cycle in these species is approximately 3 days (27). This estimate is in agreement with a previous study that obtained full blood meal microsatellite profiles 32 hours post-feeding in anopheline mosquitoes, and up to 48 hours in culicines, perhaps due to a greater initial blood meal size or a slower speed of digestion in the latter (28). In contrast the age of a sample appeared to limited impact on amplification success or likely accuracy. Among sample three age groupings (dividing into samples from 1915, 1979, or 2004 and later) amplification success was significantly different (Fisher exact test P=0.00012), but this difference was underpinned by the poor amplification of a single 1915 sample, genotype success rate for which was a significant outlier (Grubb’s test P<0.05). Excluding this sample, all loci bar one in a single 1915 sample amplified successfully. DNA degradation might also lead to loss of alleles but heterozygosity was similar among the age groups even with the outlier included (χ^2^_3_ =0.33).

Multiple feeding within one gonotrophic cycle may have significant epidemiological consequences by increasing the probability of transmission events (29,30). In vector-borne disease transmission models, a key parameter is the number of bites per unit time; this parameter is raised to square to account for acquisition of infection and subsequent onward transmission. As a result, even small changes in biting frequency may have large impacts on R0, the basic reproductive number (30,31). In the field, the frequency of multiple-feeding varies and has been shown to be reduced in southern Zambia after distribution of insecticide treated nets (29), which have become common as a result of mass distribution campaigns in Sub-Saharan Africa, since the multiple-feeding mosquitoes we identified were collected (32).

The analysis also provided intriguing historical detail; for example, one set of specimens were collected by Dr. J.W.S Macfie in 1915. Dr. Macfie was Director of the Medical Research Institute in Accra, Gold Coast (present-day Ghana) and made a number of contributions to our understanding of tropical medicine in West Africa (33,34). These specimens belonged to the genus *Mansonia*, a major nuisance biter and in some areas, perhaps including Ghana, a vector of lymphatic filariasis (35). In his collection, two mosquitoes were engorged with blood with an identical genotype (Accra_1915_2 and 3 in Table 1), indicating that they had bitten the same individual, despite being collected one month apart (Figure 1A).

Another collection from 1915 belonged to Mrs. S.L.M Connal, an entomologist at the medical research institute in Yaba, Lagos, Nigeria, who made early observations on the now text-book behaviour of *Mansonia africanus* larvae to attach to aquatic plant stems for respiration (36). Within this collection, one of the two *Mansonia* specimens examined (Figure 1B) was positive for *Plasmodium falciparum*. Although in many *Anopheles* species this could indicate a developing oocyst infection, *Mansonia* spp. do not support the development of human malaria, and so the parasites must have been in the peripheral circulation of the human on which the female had last fed. This is also very likely to be the case for the two *Anopheles* specimens, both collected from Kenya in 1979 (Table 1, Figure 1C), in which *P. falciparum* were identified (along with multiple blood-feeding in the Jimbo specimen). Though *A. quadriannulatus* is a morphologically-identical sister species of the World’s most important malaria-transmitting species, *A. gambiae*, it rarely bites humans and also displays a more efficient parasite-melanizing response to infection, effectively eliminating any significant potential for *P. falciparum* transmission (37).

## Conclusion

The ability to identify species, bloodmeal origins and infections from archived specimens stored in a manner which is far from optimal for DNA preservation opens up the possibility of mining these biobanks to better understand the evolution of vector:parasite:host systems. In museum archives, comprehensive, often well documented, entomological collections exist that pre-date anti-parasitic and anti-vector interventions, providing the additional tantalising possibility that we can use them to observe how anthropogenic forces have shaped the genomes of these organisms.

## Acknowledgements

We thank Keith Steen and Emily Rippon for assistance in the laboratory. This work was made possible by the careful curation of mosquito specimens by Dr. Evelyn Klinkenberg, Professor Harold Townson, Dr J.W.S Macfie and Mrs S.L.M. Connal.

---

**Supplementary table 1:**
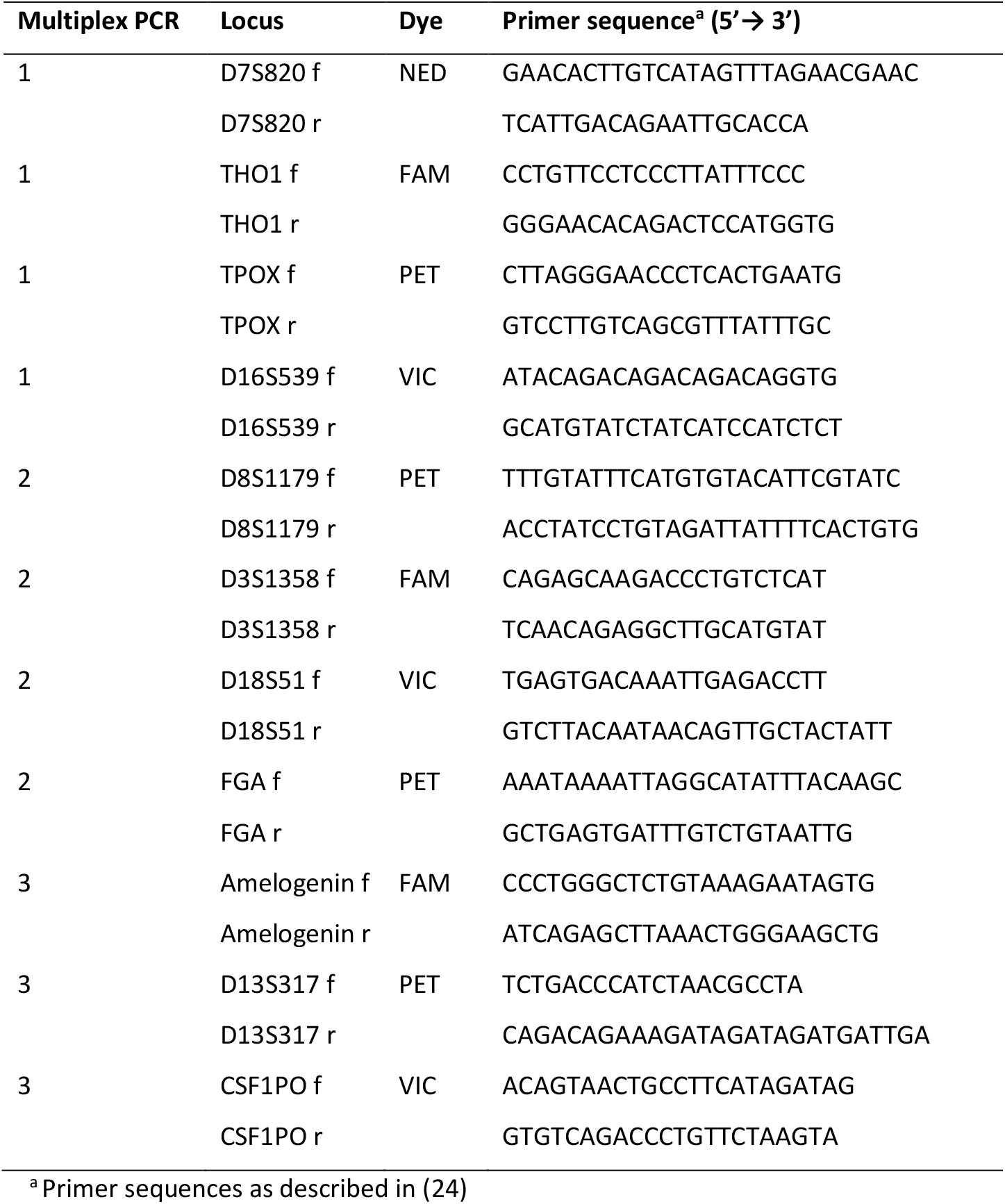
Primer sequences and fluorescence labels of the 11 microsatellite loci included in the three microsatellite multiplexes.

**Supplementary figure 1:**
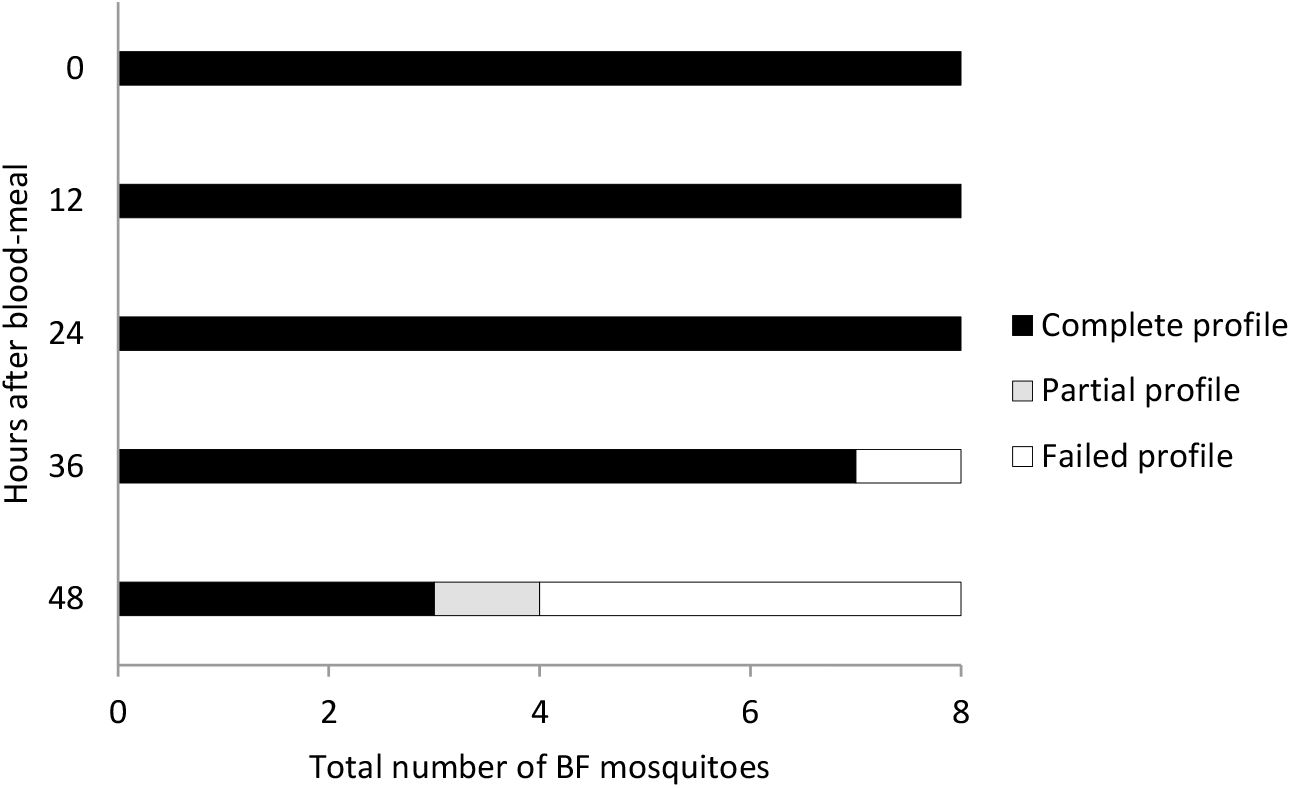
Microsatellite detection time course in *Anopheles gambiae*. Laboratory mosquitoes were killed at 5 different time points, i.e. 0h, 12h, 24h, 36h and 48h, after blood feeding. Allelic profiles were considered as complete when all 10 microsatellite loci were detected, partial if at least one locus was detected or failed when no loci were amplified.

**Supplementary figure 2:**
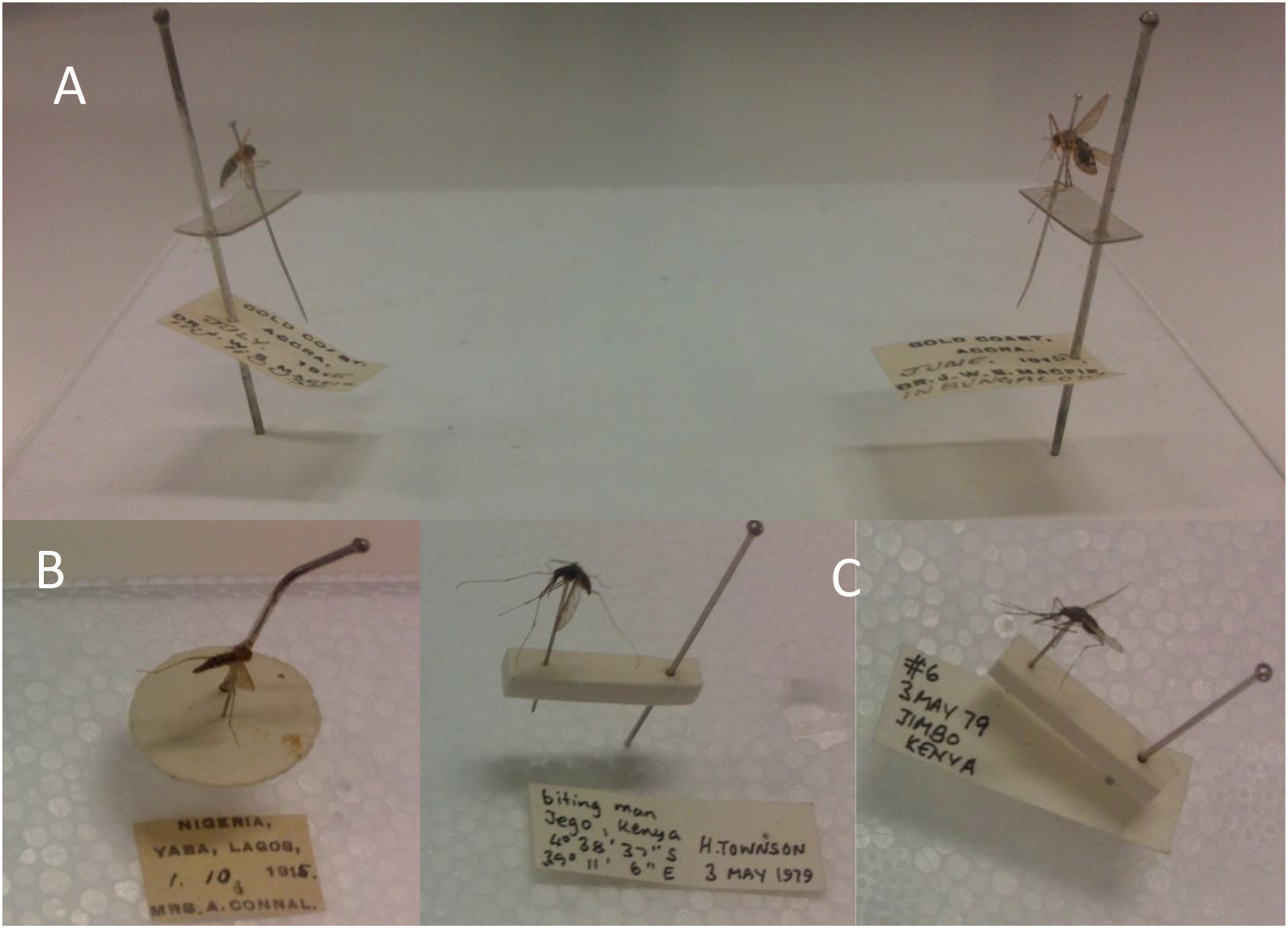
Historical mosquito collections. **A)** Two *Mansonia* specimens collected by Dr. J.W.S Macfie in Accra, Gold Coast, 1915. **B)** A *Mansonia* specimen collected by Mrs. S.L.M Connal in Yaba, Nigeria, 1915, which was positive for *Plasmodium falciparum*. **C)** Two *plasmodium*-positive *Anopheles quaddriannulatus* specimens collected by Professor Harold Townson in Kenya, 1979.

